# Tissue Registration and Exploration User Interfaces in support of a Human Reference Atlas

**DOI:** 10.1101/2021.12.30.474265

**Authors:** Katy Börner, Andreas Bueckle, Bruce W. Herr, Leonard E. Cross, Ellen M. Quardokus, Elizabeth G. Record, Yingnan Ju, Jonathan C. Silverstein, Kristen M. Browne, Sanjay Jain, Clive H. Wasserfall, Marda L. Jorgensen, Jeffrey M. Spraggins, Nathan H. Patterson, Griffin M. Weber

**Affiliations:** Department of Intelligent Systems Engineering, Luddy School of Informatics, Computing, and Engineering, Indiana University, Bloomington, Indiana, USA; Department of Biomedical Informatics, University of Pittsburgh School of Medicine, Pittsburgh, Pennsylvania, USA; Bioinformatics and Computational Biosciences Branch, Office of Cyber Infrastructure and Computational Biology, National Institute of Allergy and Infectious Diseases, National Institutes of Health, Bethesda, Maryland, USA; Department of Medicine, Washington University School of Medicine, Saint Louis, Missouri, USA; Departments of Pathology and Pediatrics, University of Florida, Gainesville, Florida, USA; Mass Spectrometry Research Center, Vanderbilt University, Nashville, Tennessee, USA; Department of Biomedical Informatics, Harvard Medical School, Boston, Massachusetts, USA

## Abstract

Several international consortia are collaborating to construct a human reference atlas, which is a comprehensive, high-resolution, three-dimensional atlas of all the cells in the healthy human body. Laboratories around the world are collecting tissue specimens from donors varying in sex, age, ethnicity, and body mass index. However, integrating and harmonizing tissue data across 20+ organs and more than 15 bulk and spatial single-cell assay types poses diverse challenges. Here we present the software tools and user interfaces developed to annotate (“register”) and explore the collected tissue data. A key part of these tools is a common coordinate framework, which provides standard terminologies and data structures for describing specimens, biological structures, and spatial positions linked to existing ontologies. As of December 2021, the “registration” user interface has been used to harmonize and make publicly available data on 6,178 tissue sections from 2,698 tissue blocks collected by the Human Biomolecular Atlas Program, the Stimulating Peripheral Activity to Relieve Conditions program, the Human Cell Atlas, the Kidney Precision Medicine Project, and the Genotype Tissue Expression project. The second “exploration” user interface enables consortia to evaluate data quality and coverage, explore tissue data in the context of the human body, and guide data acquisition.

## Main

The National Institutes of Health funded HuBMAP consortium was formed in 2018 to develop an open and global platform to map the estimated 37 trillion healthy cells in the human body^1,2^. An essential component of this platform are two software programs whose user interfaces enable organ experts to “register” tissue data and researchers to “explore” those data in a spatially and semantically explicit manner. The software programs are open and generalizable; and, they have been adopted by other international consortia mapping the human body. By combining and harmonizing datasets from these different consortia, the software is facilitating the collaborative construction of an overall human reference atlas (HRA), which will be more accurate and more complete than what is possible from any individual group.

However, constructing an HRA that captures macro to microanatomy of healthy adults down to the single-cell level poses diverse challenges^3^. First, *<u>specimen</u>* metadata collected by different laboratories for diverse donors (e.g., basic demographics such as sex, age) and using different assay types (e.g., build analyses like spatial transcriptomics^4,5^ or spatially explicit assays like CO-Detection by indEXing (CODEX)^6^ and associated data provenance) need to be harmonized so they can be cross-searched. Second, *<u>biological structure</u>* data (i.e., information on what anatomical or histological structures are present in the tissue blocks extracted from human organs for interrogation) need to be collected in a standard manner. Third, the *<u>spatial</u>* size of the tissue block, along with the position and rotation of the block in relation to reference organs positioned in a common three-dimensional (3D) reference space, need to be recorded.

To address these challenges, the registration and exploration user interfaces leverage a common coordinate framework (CCF), which is a “spatiotemporal computational framework for the management, integration, and analysis of anatomically and spatially indexed data ‘‘^7^. It creates “an underlying reference map of organs, tissues, or cells that allows new individual samples to be mapped to determine the relative location of structural regions between samples” ^8^. In 2017, NIH organized a CCF meeting^9^ which reviewed different atlas construction projects. The Allen Mouse Brain Atlas10 was highlighted as a successful example of combining large-scale mapping, quantification, analysis, and visualization of data. Brain atlas projects identify anatomic-scale landmarks (e.g., specific folds, specific lumens with cerebrospinal fluid, or major arteries) that are conserved between individuals and apply mathematical coordinate mappings to register data from individual donor tissue samples to a reference geometry, allowing direct comparison across individuals. In particular, the Talairach and the Montreal Neurological Institute Hospital (MNI) stereotaxic space emerged as CCFs for human brain tissue. The Paxinos’ Rat brain stereotaxic coordinate system and Waxholm space (WHS)^11,12^ are widely used for rodent brain^13–17^. However, a CCF that works for the brain^18^ does not necessarily work for other human organs that might be much larger (large intestine), deflated (lung), or highly variable in size (lymph nodes). We therefore constructed a CCF for the human reference atlas (CCF-HRA) which combines expert-curated ontologies for specimen, biological structure, and spatial data.

## Results

### Tissue Registration User Interface (RUI)

Groups contributing tissue data to advance the HRA need an effective means to register their data within a well-defined 3D spatial reference system and derive anatomical structure annotations. Specifically, they need to select the most appropriate reference organ (e.g., the right kidney if this is the organ from which tissue was extracted) and specify the size, position, and rotation of a tissue block in relation to that organ. They must be able to review the listing of automatically assigned anatomical structure annotations and make changes as needed (add or delete annotations) to describe what anatomical regions the tissue block captures.

The Registration User Interface (RUI) supports tissue data from specimens extracted either as tissue blocks or via biopsies. It allows users to document the tissue extraction site, in relation to a reference organ, by sizing a virtual tissue block and drag-and-drop positioning this block (in yellow in **Fig. 1a**) inside a reference organ from the 3D Reference Object Library^19^. Collision detection (the intersection of bounding volumes detected by an algorithm; see anatomical structures that make up the kidney reference in **Fig. 1b**) is used to assign anatomical structure annotations (see lower right of **Fig. 1a**). If tissue sections are cut from a tissue block, it is important to keep track of the z-stack of tissue sections and their spatial relationships to the tissue block (see **Fig. 1c**). A user can define extraction sites (see **Fig. 1d** and 3D Reference Object Extraction Sites, in Methods) and many tissue samples might use the very same site.

**Fig 1.**
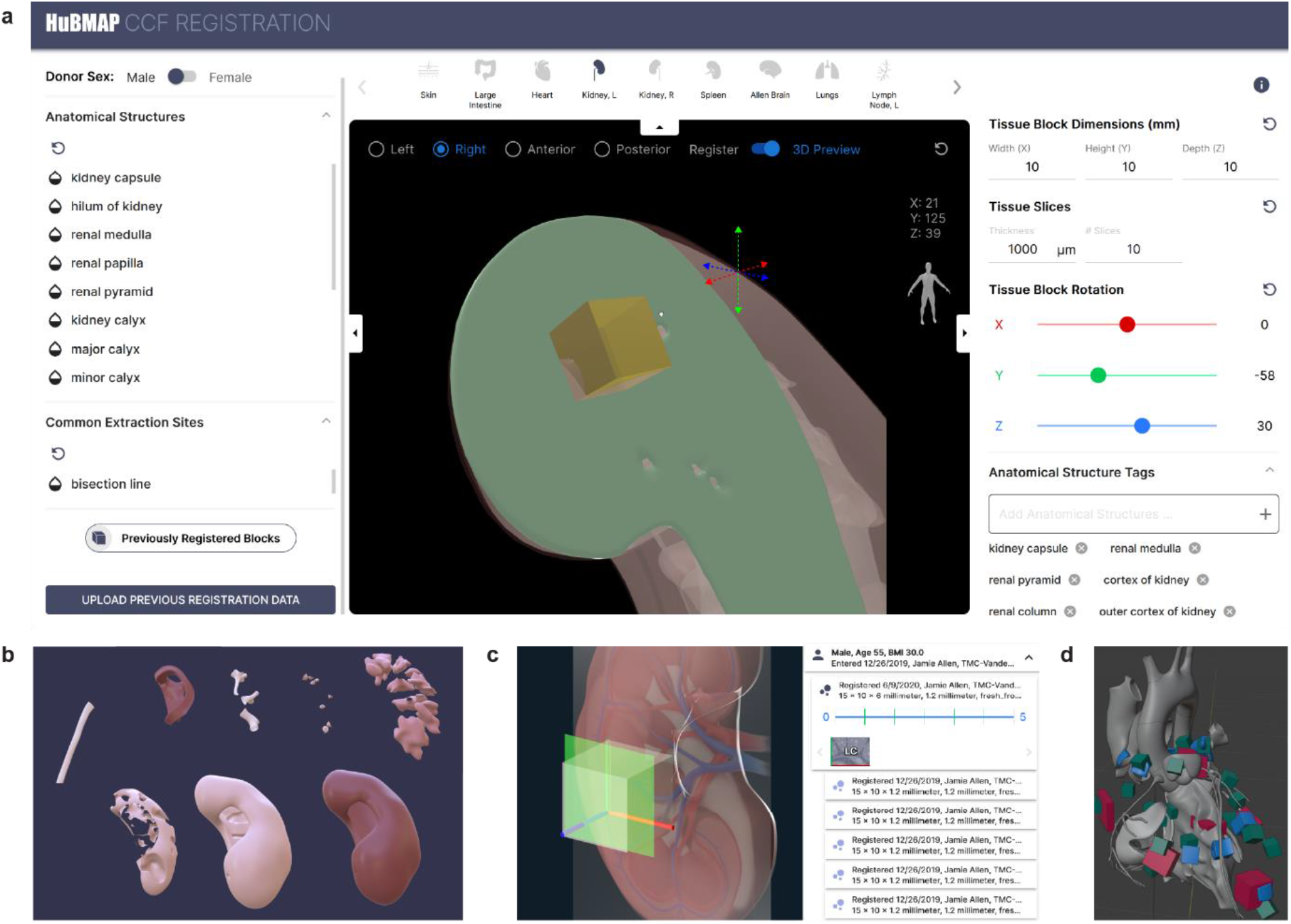
Tissue Registration User Interface (RUI). **a**. Tissue registration for kidney using the RUI. **b**. Named anatomical structures that together make up the kidney reference object. **c**. Relationship of cutting plane to x-y-z coordinate system (red, green, blue axes) of tissue block (left). Details panel on the right-hand side of EUI v2.0 with information on what tissue sections exist for a tissue block (blue lines in strip plot) and green-red axes on the left and bottom of each thumbnail. **d**. Extraction sites used by three different projects for the male heart (six sites for HCA in red, 10 sites for HuBMAP in blue, 13 sites for SPARC in green). The RUI interactive user interface is available at https://hubmapconsortium.github.io/ccf-ui/rui.

### Tissue Exploration User Interface (EUI)

The Exploration User Interface (EUI) was implemented for two stakeholders: (1) Tissue data providers can use the EUI to check and approve tissue size, location, rotation, and semantic annotations for the tissue blocks they registered with the RUI. (2) Tissue data users come to the EUI to filter, browse, and search for tissue data using specimen, anatomical structure, or spatial data queries. Users expressed interest in running queries involving anatomical structures (AS), cell types (CT), and biomarkers (B), such as: What CT are *located_in* which AS? What CT/AS are in 1mm distance from cell x? Which B’s *characterize* a certain CT in a specific AS or functional tissue unit (FTU)? How many cells of CT y are in AS x? How does a male cell census differ from a female census for the very same region (e.g., a set of AS)? Plus, there is a strong interest in running spatial queries across donors or assay types, such as: For the same 1mm^3^ in a reference body (e.g., the female body), what tissue blocks exist for a donor population (e.g., 25–30-year-old females) and/or a specific assay type (e.g., CODEX data)? Given a tissue block, what other tissue blocks exist for that same 3D volume?

The current EUI supports exploration of anatomical structures. It uses a split-screen interface (**Fig 2**): A hierarchical list of anatomical structures (e.g., “kidney: cortex”) on the left is linked to anatomically correct 3D reference objects in the middle. The data for these come from the biological structure and spatial ontologies in the CCF-HRA. A list of registered tissue blocks corresponding to the selected anatomical structure is displayed on the right, with white 3D shapes placed within the reference organs to indicate the positions of those tissue blocks. The CCF-HRA specimen ontology defines filters at the top of the user interface for patient demographics, assay type, and tissue biomarkers.

**Fig 2.**
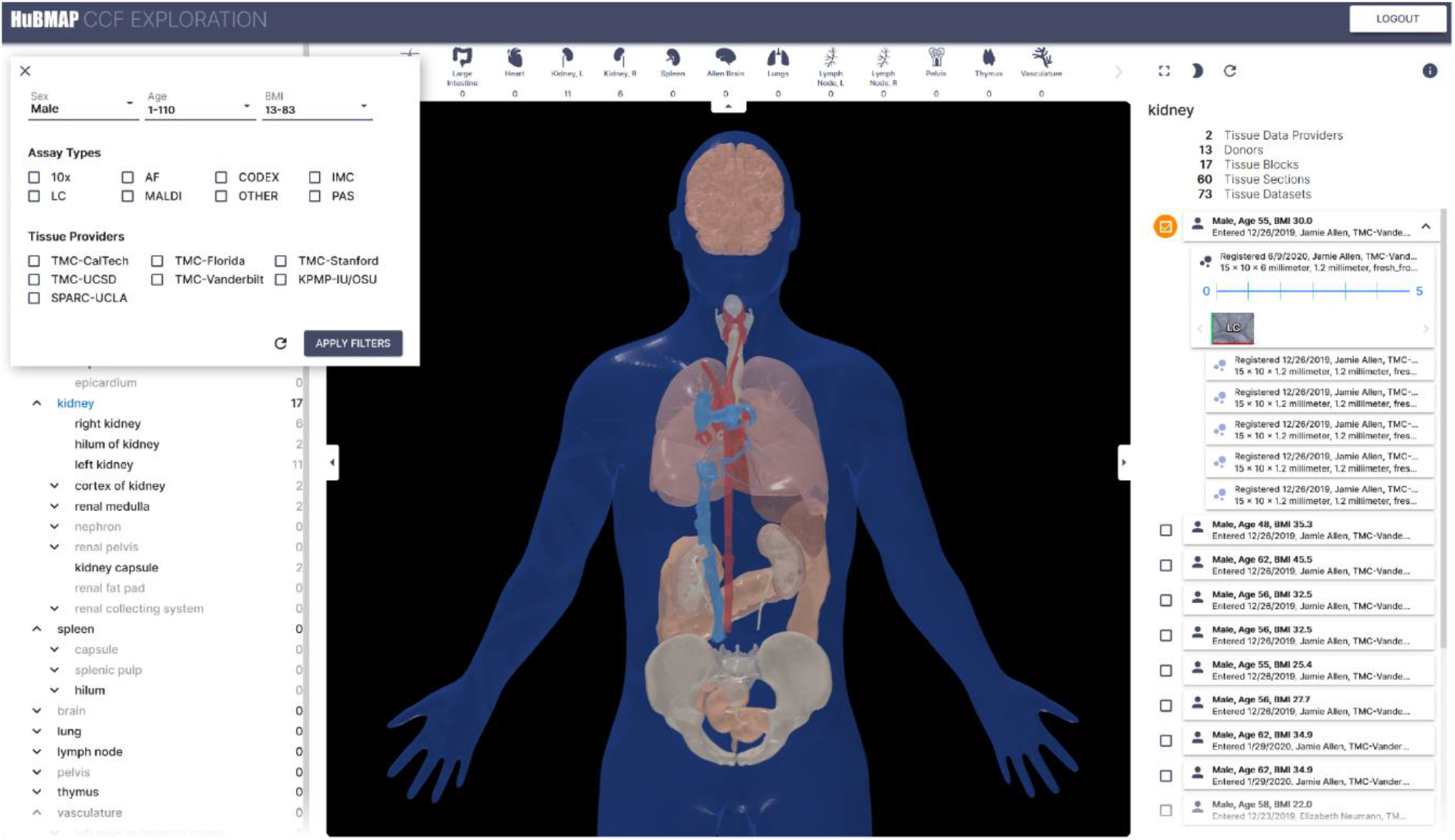
Tissue Exploration User Interface (EUI). The EUI shows the anatomical structure “partonomy” on left, 3D reference organ with registered tissue blocks (in white) in middle, and tissue sections on right. Filters for specimen metadata are provided in top middle. Registered tissue blocks can be explored via specimen data, biological structure terms, or spatially using the 3D reference organs. The EUI interactive user interface is available at https://portal.hubmapconsortium.org/ccf-eui.

### Usage of the RUI and EUI

The CCF-HRA metadata and associated 3D reference library are unique in that they interlink specimen, biological structure, and spatial data in support of human reference atlas construction and usage. The RUI and EUI use the CCF-HRA to describe collected tissue blocks and biopsies in a standardized way (**Fig 3**), which enables data harmonization across consortia, donors, organs, and assay types.

**Fig 3.**
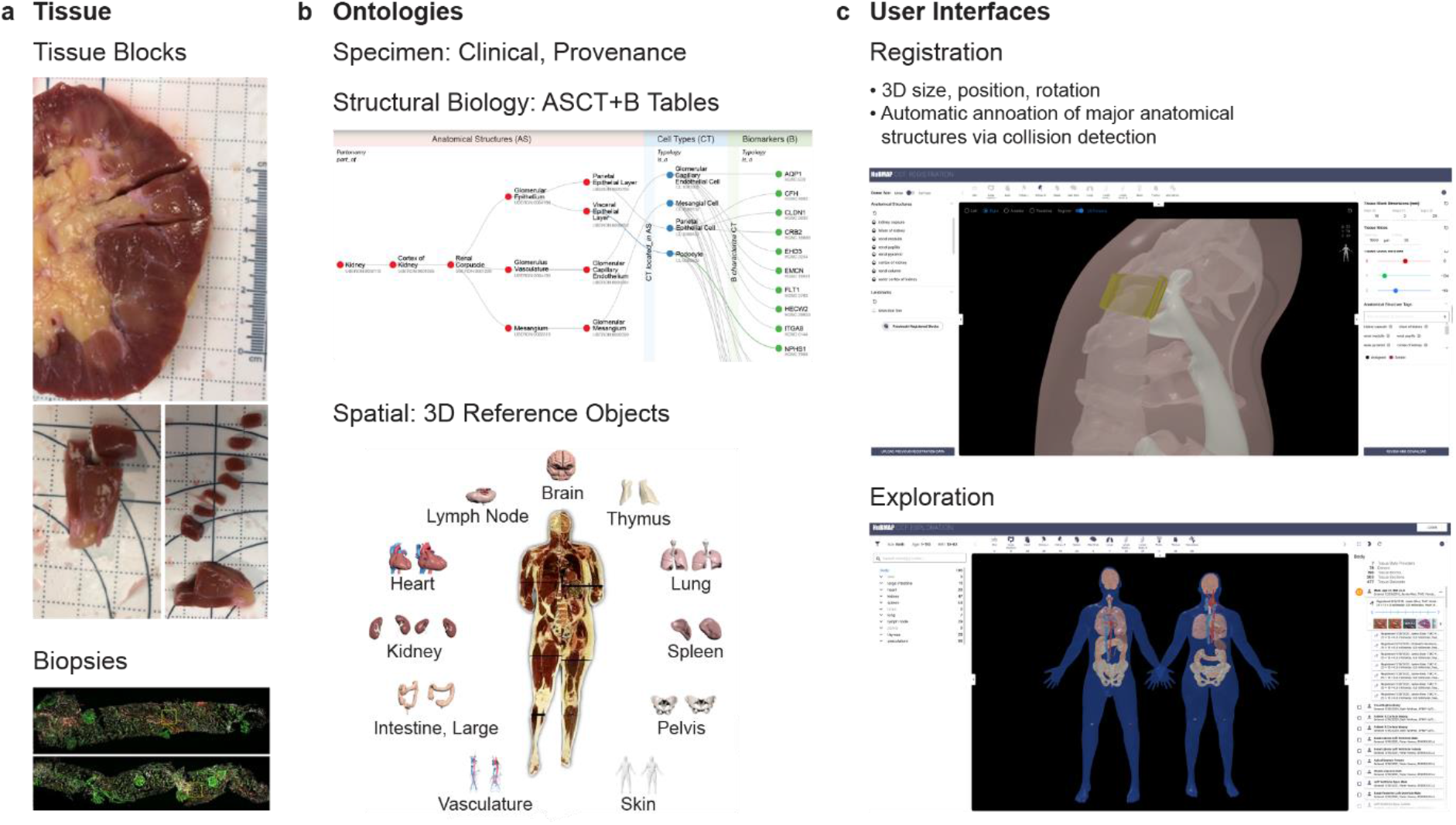
Workflow. **a**. Tissue data (blocks or biopsies) are secured by different laboratories/projects from different donors and organs. **b**. Three different ontologies capture specimen, biological structure, and spatial metadata using organ-specific ASCT+B tables and 3D reference objects. **c**. The CCF Registration User Interface (RUI) is used to register tissue data while the CCF Exploration User Interface (EUI) supports ontologically and spatially explicit exploration of tissue data across organs, donors, laboratories, and consortia.

The RUI requires about five minutes of training time and two minutes for each tissue registration^20^. As of December 2021, 404 tissue blocks and 6,178 tissue sections from the Human Biomolecular Atlas Program (HuBMAP)^3^, three biopsies from the Kidney Precision Medicine Project (KPMP)^21^, 26 heart tissue blocks from the Stimulating Peripheral Activity to Relieve Conditions (SPARC)^22^, 2,259 tissue blocks from the Genotype Tissue Expression project (GTEx)^23,24^, and six datasets from the Human Cell Atlas (HCA)^18,25,26^ have been registered. The resulting data covers 11 organs; 376 distinct data files linked to 2,698 tissue blocks and their 6,178 tissue sections.

The average size of the 404 HuBMAP tissue blocks is 15.6 mm width, 13.6 mm height, and 9 mm depth. Exactly 49 tissue blocks have associated scRNAseq gene biomarker data from which cell type annotations can be derived using Azimuth^27^ or similar services; 75 tissue blocks use validated antibody panels that detect well defined protein biomarkers and the cell types those characterize—supporting cell type identification at the single-cell level and are spatially explicit, e.g., they use CODEX^6^, Imaging Mass Spectrometry (IMS)^28^, Sequential Fluorescence In-Situ Hybridization (seqFISH)^29^, or Visium^5^, making it possible to run segmentation algorithms at the anatomical structure, FTU, or single-cell level. Examples of spatially explicit datasets are the six heart tissue blocks^30^, the 18 RUI-registered kidney tissue blocks^31^, or the 12 skin tissue blocks^32^. GTEx^24^ is an example of having multiple datasets from the same extraction sites, as defined by their standard operating procedures (SOPs)^33^.

An additional important use case for the RUI and EUI is to evaluate the coverage and quality of the HRA to inform future tissue data acquisition and analysis. Single-cell analysis is costly (e.g., scRNAseq data costs about $1 per cell; sampling a tissue block of 1mm^3^ filled with 10μm diameter cells costs about $1 million). Smart data sampling is needed to arrive at data that captures not only key anatomical structures of single organs but also the diversity of healthy male and female adults. The CCF-HRA and the presented user interfaces make it possible to compare the coverage of data across projects, as illustrated in the HRA Dashboard (**Supplementary Figure 1**).

## Discussion

The RUI and EUI were released in May 2021. Since then, they have been adopted by five consortia contributing data towards the HRA. Both user interfaces are available as web components, which have been successfully integrated in the HuBMAP portal and will soon become available in the GTEx portal. The CCF-HRA’s three ontologies have been fully implemented for eleven organs. They enable the RUI and EUI to capture (1) specimen metadata about the donor (the “who”); (2) biological structure data, which describes “what” part of the body a tissue sample came from and details anatomical structures, cell types, and biomarkers of the tissue sample; and (3) spatial data, which indicates “where” a tissue sample is located in a shared 3D common coordinate system.

### Limitations

Known limitations for the RUI include a limited set of reference organs, limited anatomical structures (small structures are difficult to model in 3D), and scaling tissue blocks based on the relationship between the original organ size and the reference organ size. Additionally, manipulating the 3D position and rotation of virtual tissue blocks in a 3D scene inside a browser can be challenging, especially for users with little or no experience with 3D software. However, recent work has shown that position accuracy, rotation accuracy, and completion is still achievable with some training^20^.

A user study that examines task accuracy and completion time for different EUI exploration tasks (e.g., those discussed in the User Requirements section) is in progress. Known limitations for the EUI include limited spatial querying, limited filters, and no deep linking facilities. Further, the 3D navigation of the organs in their spatial context can be challenging for novice users. Finally, extended loading times may be possible (depending on the browser).

While the CCF-HRA is a work-in-progress, the current data structures and user interfaces demonstrate the entire workflow from human tissue acquisition and registration via the CCF RUI to data representation and exploration within the CCF EUI for a rigorously defined data registration and management process. We are sharing this first CCF-HRA implementation to build awareness of the work we are doing and to obtain feedback and suggestions from the broader community, including other efforts to map the human body at single-cell resolution. In addition, we seek feedback from scientists who are interested in using HuBMAP data for their research, integrating CCF-HRA user interfaces into their workflows, and/or incorporating CCF-HRA data structures and user interfaces into their portals.

### Next Steps

The ultimate goal of this effort is to develop effective means to interlink and federate new and existing data across scales (anatomical structures, cell types, and biomarkers) and to link the evolving HRA to existing ontologies and 3D references. This will make it possible to understand and communicate the spatial organization of the human body across scales; interpret new datasets in the context of the evolving CCF-HRA (e.g., support CT label transfer as done by Azimuth^27^); and map tissue blocks spatially based on their unique morphology and molecular “fingerprint” (e.g., given scRNA-seq data for a new tissue block, this block can be mapped into the 3D space that has a similar cell-type population with comparable biomolecular states).

New datasets, technologies, and user requirements will both demand and make possible continuous improvements of the 3D reference object library and ontologies, the registration, mapping, and exploration processes, and the user interfaces. The CCF-HRA and associated UIs are expected to evolve to support ever more robust and detailed registration and exploration of semantically and spatially annotated tissue data.

Developing a CCF-HRA for the human body is a major undertaking that requires access not only to high-quality and high-coverage data but also to human expertise across both biological domains and technological domains. It seems highly desirable to develop and agree on data formats across consortia and to develop data infrastructures and user interfaces that empower many users to contribute to the construction and usage of a human reference atlas. Serving as a “rosetta stone,” the ASCT+B tables help translate and federate data across organs and make it possible to have many experts contribute. User interfaces like the CCF RUI and EUI make it possible to register tissue samples spatially and semantically in a uniform manner in support of cross-consortium exploration via the EUI.

Experts interested in joining this international effort are invited to register at https://iu.co1.qualtrics.com/jfe/form/SV_bpaBhIr8XfdiNRH to receive regular updates and invites to relevant meetings that aim to advance the construction and usage of a human reference atlas.

## Methods

### CCF-HRA Specimen Ontology and Data

CCF-HRA Specimen ontology data describe the demographic, clinical, and other metadata associated with human specimen tissue samples. The complete HuBMAP clinical data— covering more than 100 metadata fields—was reduced to a smaller set of metadata fields that are relevant for CCF-HRA construction and usage. CCF-HRA Specimen data was then compared with data made available by other consortia to finalize data fields relevant for CCF-HRA design. The current CCF-HRA specimen data includes demographics and clinical data (e.g., sex, age, body mass index [BMI]), workflow information (e.g., tissue sample creation/modification date, donor/organ/tissue ID, specimen/data/assay type), author information (e.g., author group/creator), and links to source data. Additional components of the CCF-HRA Specimen data link samples to the laboratories (tissue mapping centers [TMCs] in HuBMAP) and projects Kidney Precision Medicine Project (KPMP)^34^, Stimulating Peripheral Activity to Relieve Conditions (SPARC)^22,35^, Genotype-Tissue Expression (GTEx)^23^ that collected the tissue, and assay types (e.g., CODEX, Periodic acid–Schiff (PAS), or autofluorescence (AF). Note that for HuBMAP data, specimen information is created on the fly by querying the HuBMAP search-API. For data from other consortia, specimen information is stored in the *rui_locations*.*jsonld* for each tissue block.

The complete set of specimen data consist of data collected during data ingestion and data generated during tissue registration using the RUI. Data is shared in JSON-LD with the following 22 fields: donor ID (de-identified), age (or age range), sex, BMI (kg/m^2^), consortium name, tissue provider name, tissue provider UUID, tissue provider author, links to source data, tissue block ID, tissue block dimensions (width, height, depth, and units of measure), tissue block position (relative to a CCF-HRA reference organ), tissue block rotation (relative to a CCF-HRA reference organ), tissue block placement author, number of tissue sections (for a given tissue block), size of tissue sections (including units of measure), tissue section ID, tissue section number, dataset ID, dataset technology used, dataset assay type, and dataset thumbnail.

### CCF-HRA Biological Structure Ontology and Data

In addition to registration and exploration user interfaces, the user requirements assessment identified the need for agreement across multiple consortia and stakeholders on what anatomical structures (AS), cell types (CT), and biomarkers (B) are relevant for construction of an HRA. The biomarkers include cell type-specific genes, proteins, lipids, and metabolites commonly used to characterize these CTs. Collectively, we call this the ASCT+B terminology. Furthermore, anatomical structures and cells exist in a 3D context, and there is a need for 3D representations of major AS and CT so that experimental data can be spatially registered, explored, and analyzed in the context of this 3D HRA.

A parallel effort by 16 consortia^36^ is therefore creating (1) ASCT+B data tables that capture major AS, CT, and B and their interrelationships—including *part_of* relations (the hierarchical “partonomy”) between AS, *located_in* relations between CT and AS, and *characterizes* relationships between B and CT and (2) associated reference objects that represent the 3D size, position, rotation, and shape of major AS that are listed in the ASCT+B tables. While organs differ substantially in their form and function and across human specimens, all have AS, CT, and B that are captured in the ASCT+B tables and the associated 3D Reference Object Library^37^. The resulting ASCT+B tables represent expert knowledge and information from textbooks, scholarly publications, and experimental data evidence. An associated 3D Reference Object Library captures the anatomically correct 3D shapes, sizes, locations, and rotations of key AS present in the ASCT+B tables. The organs are developed by specialist 3D medical illustrators, and they are approved by organ experts^38,39^. See **Fig 4a** for an exemplary partial ASCT+B table and corresponding 3D Reference Objects of a kidney.

**Fig 4.**
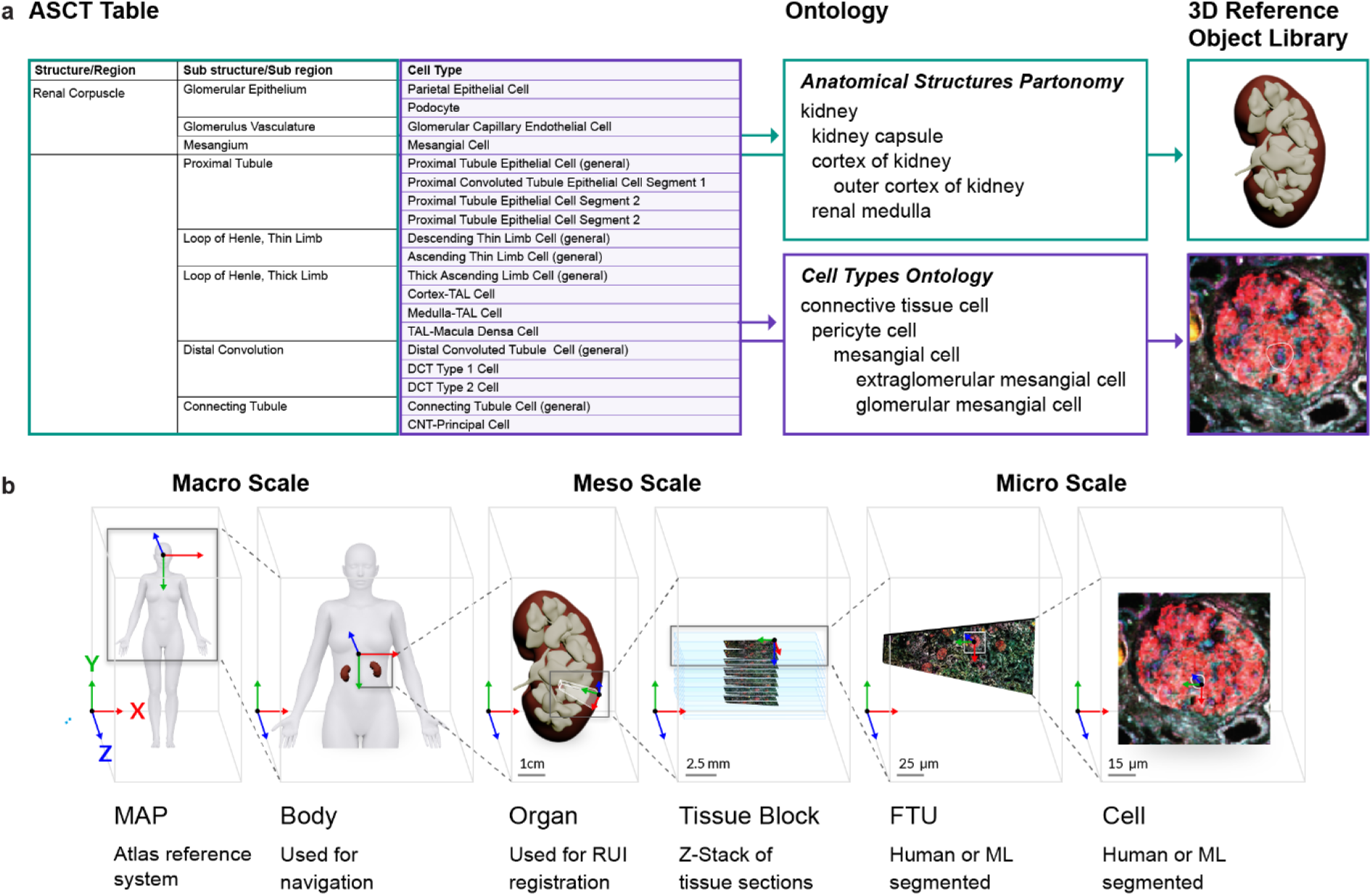
Semantic and Spatial Representation of a Kidney. **a**. The CCF Biological Structure Ontology represents the body as a set of nested, named anatomical structures (the AS “partonomy”) and cell types (the CT “typology”, not shown). AS and CT are linked to Uberon/FMA and CL ontologies, and are mapped to 3D Reference Objects at macro to single-cell level. **b**. The CCF Spatial Ontology leverages a 3D Reference Object Library to define the dimensions and shapes of ASCT+B entities within the HRA 3D space (called the Atlas reference system). The male and female VHP reference bodies are registered within the HRA space. All organ reference objects are registered to the respective male or female VHP reference body. A new tissue block is sized, positioned, and rotated relative to a reference organ. Segmentation masks for single cells and functional tissue units identified via manual or automatic means are spatially registered with their respective tissue images (z-stacks of 2D tissue sections make up 3D tissue blocks).

The standard operating procedure (SOP) for ASCT+B table construction details this process^40^. In short, AS, CT, and B names are compiled with their unique identifiers in existing ontologies, such as Foundational Model of Anatomy (FMA)^41–43^, Uber-anatomy ontology (Uberon)^44^, Cell Ontology^45^, and Human Genome Nomenclature Committee (HGNC)^46^ in machine readable, computable format. Note that the existing ontologies have tens of thousands of terms, many of which are not relevant for healthy human adults (e.g., terms for capturing development and growth, cross-species comparisons, and disease). By authoring the ASCT+B tables, we have discovered omissions in existing ontologies that need to be resolved to properly represent healthy human adults. In these cases, we are working closely with the respective ontology curators to add these terms and corresponding literature justifications to the ontologies so they properly represent the human reference atlas. The resulting CCF-HRA Biological Structure exclusively captures terms relevant for healthy human adults. An ASCT+B table Reporter website was built to visualize the CCF-HRA biological structure data (**Supplementary Figure 2**). It facilitates the construction of the ASCT+B tables by domain experts and enables new users of the RUI and EUI to explore and become familiar with the data.

In March 2021, 11 ASCT+B tables and 26 associated reference organs were published, Digital Object Identifiers (DOIs) were minted, and all files were made publicly available^47^. In addition, the ASCT+B tables were published using Semantic Web technologies in Web Ontology Language (OWL 2)^48^, (see details of the CCF-HRA Model in Methods section). The CCF.OWL was deposited in BioPortal^49^ which supports version control. Counts of the number of entities and relationships for the 11 organs can be found in **Supplementary Table 1**. For example, the ASCT+B for the kidney features 61 AS, 62 CT, and 150 gene biomarkers (BG), but also 62 *part_of* relationships between AS, 60 *located_in* relationships from CT to AS, and 257 *characterize* relationships from BG to CT.

Using the CCF.OWL, the 3D reference objects of the HRA (but also tissue blocks registered into them) are compatible with and can be linked to other ontologies and the data they describe. For example, anatomical structures or genes can be linked to chemical compounds in the ChEMBL^50–52^, Chemical Entities of Biological Interest (ChEBI)^53,54^ or diseases using the Disease Ontology^55^ and Human Phenotype Ontology (HPO)^56–60^.

### CCF-HRA Spatial Ontology and Data

CCF-HRA Spatial data describes the 2D and 3D shapes of entities and their physical size, shape, location, and rotation—from the whole-body level down to single cells (**Fig 4b**). There are three key concepts to make this work: (1) A *Spatial Entity* represents a real-world thing (e.g., a human body, a human kidney, a tissue block or section, or an individual cell). It defines a bounded Cartesian space and its measurement units. By using the *ccf_representation_of* property, we say that a *Spatial Entity* is representing/standing in for either an ASCT+B term in the CCF-HRA Biological Structure or a physical object, such as a tissue sample. *Spatial Entities* connect to ASCT+B terms using either *ccf_representation_of* or *ccf_annotation*. (2) A *Spatial Object Reference* points to an object in the CCF-HRA Reference Object Library^37^. It provides a reference to an external representation of a *Spatial Entity*, such as a 3D object file (e.g., in obj, fbx, gltf format) or a 2D image (e.g., in tiff, png, svg format). Initially, most of the anatomically correct 3D reference organs in this library are created using male and female data from the Visible Human Project made available by the National Library of Medicine^61–63^. (3) A *Spatial Placement* defines how to place a *Spatial Entity* or *Spatial Object Reference* relative to another *Spatial Entity* using scaling, rotation, and translation (in that order). Note that rotation (in x, y, z order) occurs around the center of the object’s coordinate space; by default, rotation is considered in Euler order. In the case of *Spatial Object References*, it defines how to transform a 2D or 3D object so that it fits the *Spatial Entity*’s dimensions and units. In the case of *Spatial Entities*, it shows how to place one *Spatial Entity* relative to another.

A unique feature of the CCF-HRA is the interlinkage of the ASCT+B to the 3D-Reference-Organs. A Crosswalk table^64^ links AS in the ASCT+B tables to AS in the 3D organ file. Given this interlinkage of ontology terms and 3D nested objects, new forms of semantic and spatial queries can be supported. For example, given a registered tissue block, the 3D AS it contains/collides with can be determined and used as annotations; using these AS annotations, all AS and CT inside of these can be retrieved. The crosswalked data also makes it possible to use CT as search queries and to retrieve all AS that typically contain these CT.

3D object scene graphs are created using spatial entities, spatial object references, and spatial placements. Spatial entities and spatial object references are placed relative to other spatial entities to create a 3D scene graph.

In addition to the 3D Reference Objects, 3D spatial entities are created for extraction sites and tissue blocks. In general, each tissue block has a well-defined extraction site in relation to a well-defined reference object. In addition, we capture segmentation masks for functional tissue units. This data and their relationships are captured using a standard JSON-LD data format.

#### 3D Extraction Sites and Landmarks

In support of systematic tissue extraction across human donors, many laboratories/projects sample tissue using well defined “extraction sites.” Each extraction site defines the size, location, and rotation of a tissue block in relation to a reference organ, along with a set of anatomical structure annotations derived via collision detection (see sample in **Fig. 3a**). Given a new tissue sample, the previously defined extraction site is used as metadata. The RUI makes it easy to define an extraction site. For example, for the heart, 6 extraction sites have been identified for HCA, 10 for HuBMAP, 13 for SPARC (see **Fig. 3b**), and 14 for GTEx. The process of defining extraction sites is captured in an SOP^65^. Named landmarks are available in GLB and FBX format and are saved as female and male reference organs^66^. There are four large intestines, one kidney, and three spleen extraction sites. Further, the GLB files are semantically tagged and linked into the CCF-HRA Ontology (CCF.OWL) via the JSON-LD data.

#### 3D Tissue Blocks

Using the RUI, metadata can be generated for tissue blocks. All data can be inspected, corrected, and downloaded in JSON-LD format (see example data file in **Supplementary Figure 3**). Note that anatomical structure annotations ‘*ccf_annotation*’ are provided as PURL links to Uberon terms. ‘Scaling’ indicates how much to shrink or expand a tissue block and is defined as a ratio. ‘Translation’ captures how to translate a *SpatialEntity* (e.g., a tissue block) so that it is placed relative to another *SpatialEntity* (e.g., a reference organ) and it is rounded to 1nm precision. In some cases, tissue sections are cut from a tissue block. RUI users can enter information on tissue section thickness under ‘*slice_thickness*,’ which is given in micrometers. The number of RUI locations publicly available for a HuBMAP tissue block can be accessed in the HRA Dashboard (**Supplementary Figure 1**).

#### 3D Segmentation Masks

We distinguish three types of annotated segmentation masks: anatomical structure (AS), Functional Tissue Unit (FTU), and single-cell (SC) masks. Exemplary masks were compiled for and used in the 1st HuBMAP Kaggle competition^67^. The AS and FTU masks are modified GeoJSON files that capture the outline of annotations by their pixel coordinates, and they were generated from annotations by subject matter experts using QuPath^68^. **Supplementary Figure 4** shows a JSON file that describes one glomeruli mask for a kidney. The yellow highlighted part lists the x, y locations of the 15 line segments that comprise the polyline that outlines one glomerulus shown on the right.

#### Biological Structure Annotation via Collision Detection

When a tissue sample, such as a kidney tissue block, is registered and annotated through the RUI, it is assigned unique identifiers (e.g., “UUID-S-5678” linked to “Donor UUID-D-1234”) using the CCF-HRA Specimen Ontology. It is linked to a term in the CCF-HRA Biological Structure Ontology, indicating the anatomical structure or cell type (e.g., “kidney cortex”). The CCF-HRA Biological Structure Ontology’s anatomical structures partonomy details how the sample fits within larger structures, up to the whole body. Using the CCF-HRA Spatial Ontology, samples are linked to a *Spatial Entity* (e.g., “UUID-SE-9123”), which gives its size/dimensions. A *Spatial Placement* (e.g., “UUID-SP-4567”) positions the sample relative to another *Spatial Entity* (e.g., “#VHKidney”). After proper registration, all colliding objects are identified and corresponding AS terms from the ASCT+B tables are assigned to the tissue block as “ccf_annotations.” Further, experts are able to verify and add or remove AS terms while registering.

## Data availability

All data is open and free to access.

The ASCT+B tables and associated 3D reference objects for 11 organs are available at the CCF-HRA Portal, https://hubmapconsortium.github.io/ccf.

Registration data for HuBMAP, SPARC, HCA, KPMP, and GTEx are available at https://portal.hubmapconsortium.org/ccf-eui. For HuBMAP data, tissue blocks link to sample and donor metadata and raw data. For all other data, the blocks link to relevant papers or datasets in the SPARC, HCA, KPMP, and GTEx data portals.

All existing ASCT+B tables can be downloaded from https://hubmapconsortium.github.io/ccf/pages/ccf-anatomical-structures.html.

Organs in the CCF-HRA 3D Reference Object Library are available as GLB files at https://hubmapconsortium.github.io/ccf/pages/ccf-3d-reference-library.html. Further, these GLB files are semantically tagged and linked into the CCF-HRA Ontology (CCF.OWL) via the JSON-LD data format.

## Code availability

The registration and exploration user interfaces, standard operating procedures, and video demos of proper usage of both user interfaces are available via the CCF-HRA Portal, https://hubmapconsortium.github.io/ccf.

Code is publicly hosted at GitHub https://github.com/hubmapconsortium/ccf-ui. The Python script used to compute sizes of the 404 HuBMAP tissue blocks is at https://github.com/cns-iu/HRA-supporting-information.

The RUI is available via the HuBMAP Ingest Portal at https://portal.hubmapconsortium.org and as a stand-alone^69^ web application. A RUI video tutorial v2.3.1 is at https://youtu.be/v9ht5S7MVc0. An SOP for the RUI is available^19^.

Standard operating procedures, Demonstrations, learning modules, and self quizzes exist in the Visible Human MOOC at https://expand.iu.edu/browse/sice/cns/courses/hubmap-visible-human-mooc.

The CCF-HRA v1.6 source code repository is available at http://purl.org/ccf/source. The CCF-HRA v1.6 consists of three main parts: (1) The *CCF-HRA Core Model* is an architecture for modeling 2D and 3D spatial data annotated with ontology terms. (2) The *CCF-HRA Spatial Reference System* is a curated and annotated instantiation of the Core Model covering the entire scale from the human body down to the single-cell level. (3) The *CCF-HRA Anatomical Structures Partonomy* is a curated partonomy using existing ontology terms which mirrors the CCF-HRA Spatial Reference System.

The CCF-HRA ontology has been defined as a formal ontology. Data is added as RDF/XML or JSON-LD that is translated to RDF/XML.

## Acknowledgements

We appreciate close collaboration with fellow HuBMAP components TMC-Vanderbilt, TMC-UCSD, TMC-Florida, and the IEC, as well as with the NIH’s National Institute of Allergy and Infectious Diseases (NIAID). Bill Shirey (University of Pittsburgh) supported the users of our interfaces during the data ingest into the HuBMAP ecosystem. Kristin Ardlie (Broad Institute) facilitated data interoperability between the Genotype-Tissue Expression (GTEx) project and HuBMAP. Seth Winfree (University of Nebraska Medical Center) and Peter Hanna (University of California, Los Angeles) used the CCF RUI to register tissue blocks for the Kidney Precision Medicine Project (KPMP) and the Stimulating Peripheral Activity to Relieve Conditions (SPARC) project, respectively. Michela Noseda (Imperial College London) assisted us with spatially registering Human Cell Atlas (HCA) extraction sites and landmarks for the heart. David Osumi-Sutherland (European Molecular Biology Laboratory) and Alexander Diehl (University of Buffalo) provided expert input on extracting a non-developmental, healthy human subset of Uberon and Cell Ontology. Zorina S. Galis and Tyler Best (NIH) expertly commented on an early draft of this paper. Todd Theriault copy-edited the manuscript. Tracey Theriault assisted with the figures.

This research has been funded by the National Institutes of Health under Human Biomolecular Atlas Program (HuBMAP) awards OT2OD026671 [KB, AB, BWH, LEC, EMQ, EGR, YJ, GMW], OT2OD026675 [JCS], U54AI142766 [CHW, MLJ], U54DK120058 [JMS, NHP], and U54HL145608 [SJ]; by the Cellular Senescence Network (SenNet) Consortium Organization and Data Coordinating Center (CODCC) award 1U24CA268108-01 [KB, AB, BWH, EMQ, EGR, YJ, JCS, GMW]; by the National Institute of Diabetes and Digestive and Kidney Diseases (NIDDK) Kidney Precision Medicine Project grant U2CDK114886 [KB, EMQ, YJ]; by the Common Fund Data Ecosystem (CFDE) award OTA 20-005 OT2 OD030545 [KB, YJ]; and the National Institute of Allergy and Infectious Diseases (NIAID), Department of Health and Human Services under BCBB Support Services Contract HHSN316201300006W/HHSN27200002 [KMB]. The funders had no role in study design, data collection and analysis, decision to publish, or preparation of the manuscript. The views and conclusions contained in this document are those of the authors and should not be interpreted as representing the official policies, either expressed or implied, of the NIH.

## Author contributions

KB served as lead author and shares corresponding authorship with AB. AB, BWH, EMQ, and GMW reviewed and edited the manuscript. KB, BWH, LEC, and AB developed the RUI. AB and LEC conducted formal and informal user studies to improve usability and test new features for the RUI. BWH led the development effort for all CCF user interfaces. EGR oversaw the development and documentation of standard operating procedures. YJ worked on the segmentation masks data format with JMS, NHP, and the HuBMAP GE team. JCS and his team integrated the CCF user interfaces into the HuBMAP portal. KMB developed the 3D reference organs. SJ, CHW, MLJ, JMS, NHP authored the initial ASCT+B tables and associated 3D reference organs; they provided expert comments that helped optimize the CCF user interfaces.

## Competing interests

The authors declare no competing interests.

## Supplementary Information

**Supplementary Figure 1.**
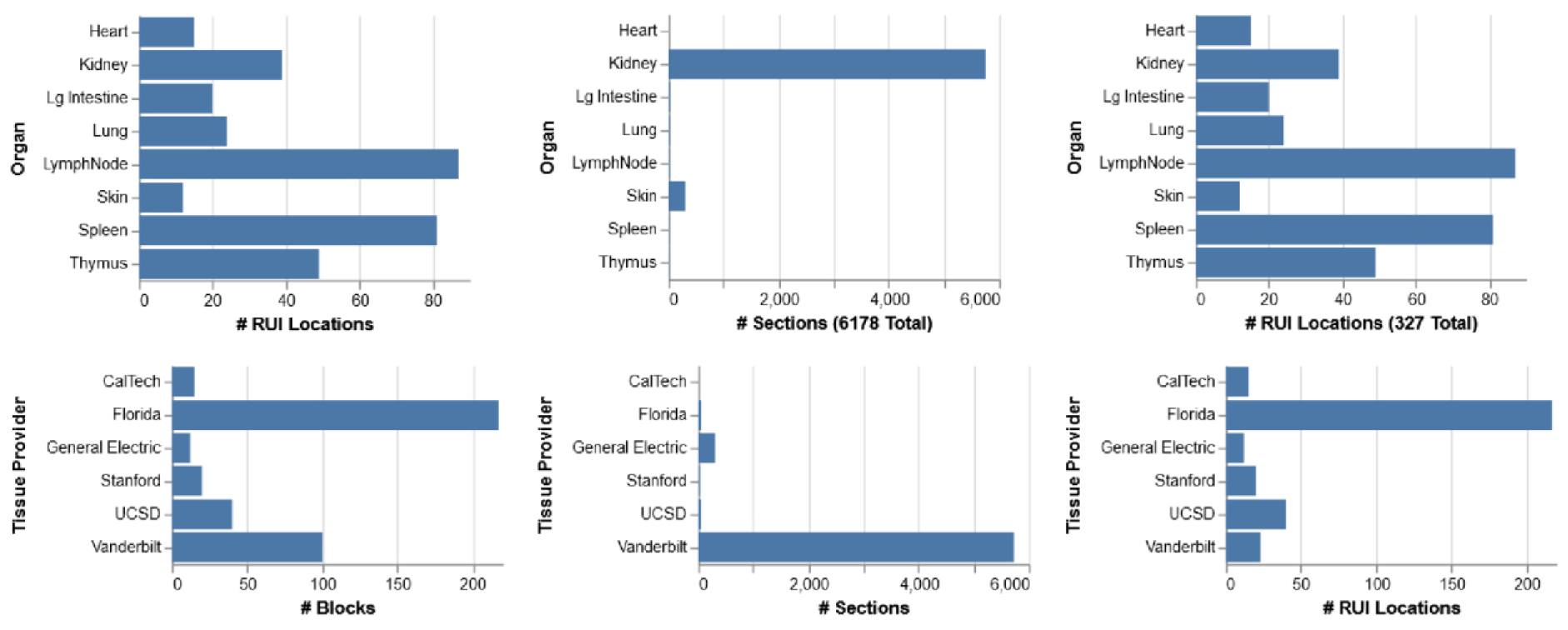
HRA Dashboard. Number of tissue blocks, tissue sections, and RUI locations per organ and tissue provider for HuBMAP data. Interactive visualization of this dashboard for all public HuBMAP data can be explored at https://hubmapconsortium.github.io/hra-data-dashboard. Note that EUI counts are higher as data by SPARC, GTEx, and other consortia are included.

**Supplementary Figure 2.**
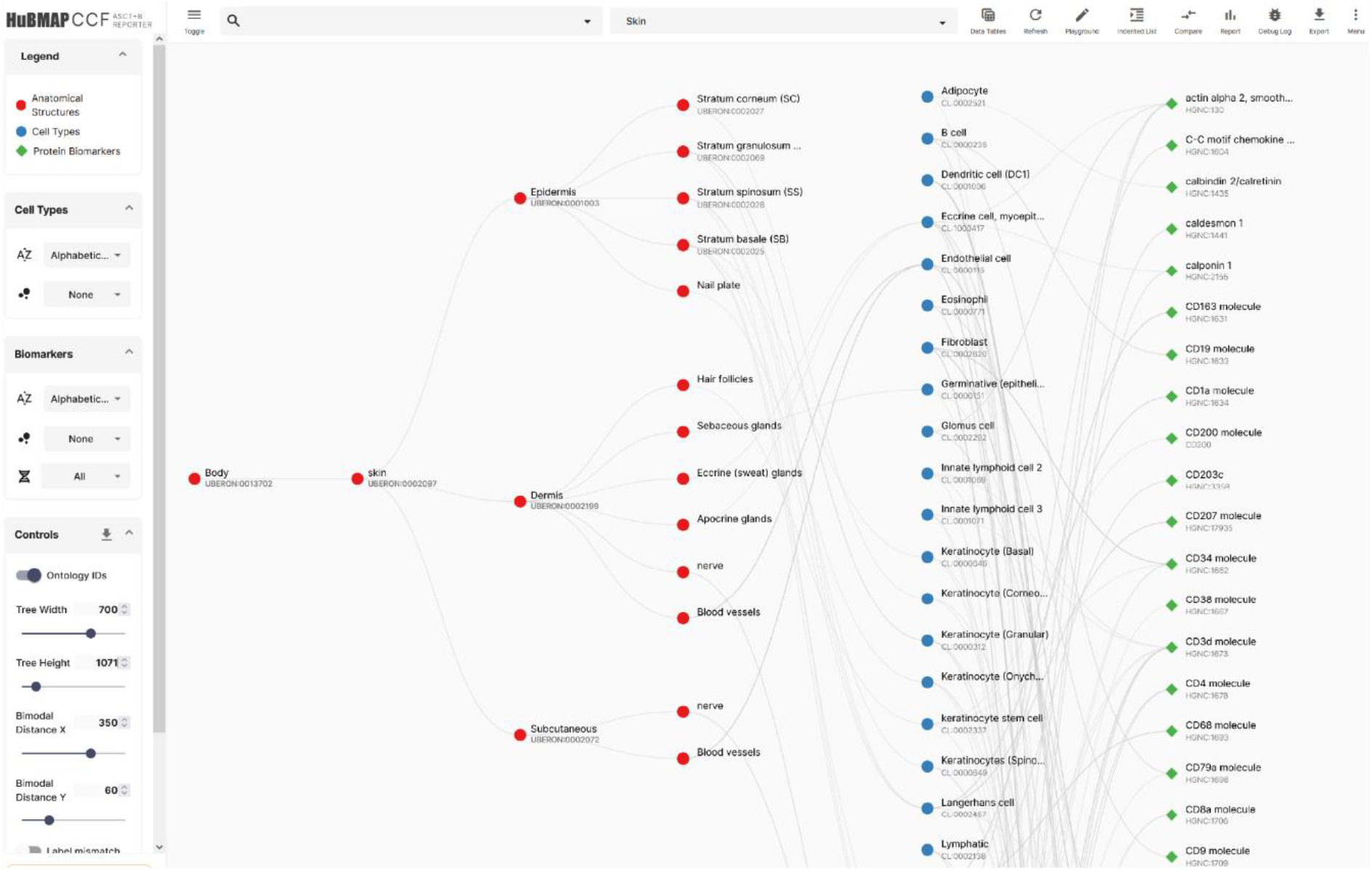
Visualizing ASCT+B Table Data. The kidney ASCT+B table captures anatomical structures (red nodes), cell types (blue), and biomarkers (green) and the interlinkages. Users can hover over a node to see details and other nodes linked to it.

**Supplementary Table 1.** ASCT+B Table Counts. Counts for anatomical structures (AS), cell-types (CT), and biomarkers (B) and their relationships for the 11 ASCT+B tables published in March 2021.

| Organ | #AS | #CT | #B Total | #BG | #BP | #AS-AS | #AS-CT | #CT-B |
| --- | --- | --- | --- | --- | --- | --- | --- | --- |
| Bone Marrow & Blood/Pelvis | 2 | 45 | 231 | 177 | 54 | 2 | 70 | 631 |
| Brain | 184 | 127 | 257 | 257 | 0 | 185 | 127 | 346 |
| Heart | 50 | 23 | 45 | 45 | 0 | 60 | 183 | 74 |
| Intestine, Large | 48 | 57 | 85 | 84 | 1 | 290 | 1,158 | 178 |
| Kidney | 61 | 62 | 150 | 150 | 0 | 62 | 60 | 257 |
| Lung | 146 | 87 | 258 | 258 | 0 | 911 | 1,658 | 722 |
| Lymph Node | 35 | 45 | 167 | 94 | 73 | 57 | 121 | 417 |
| Skin | 15 | 36 | 68 | 0 | 68 | 17 | 19 | 96 |
| Spleen | 27 | 66 | 163 | 72 | 91 | 55 | 413 | 372 |
| Thymus | 17 | 41 | 442 | 378 | 64 | 37 | 198 | 613 |
| Vasculature | 839 | 2 | 1 | 1 | 0 | 867 | 604 | 2 |
| Totals: | 1,424 | 591 | 1,867 | 1,516 | 351 | 2,543 | 4,611 | 3,708 |

**Supplementary Figure 3.**
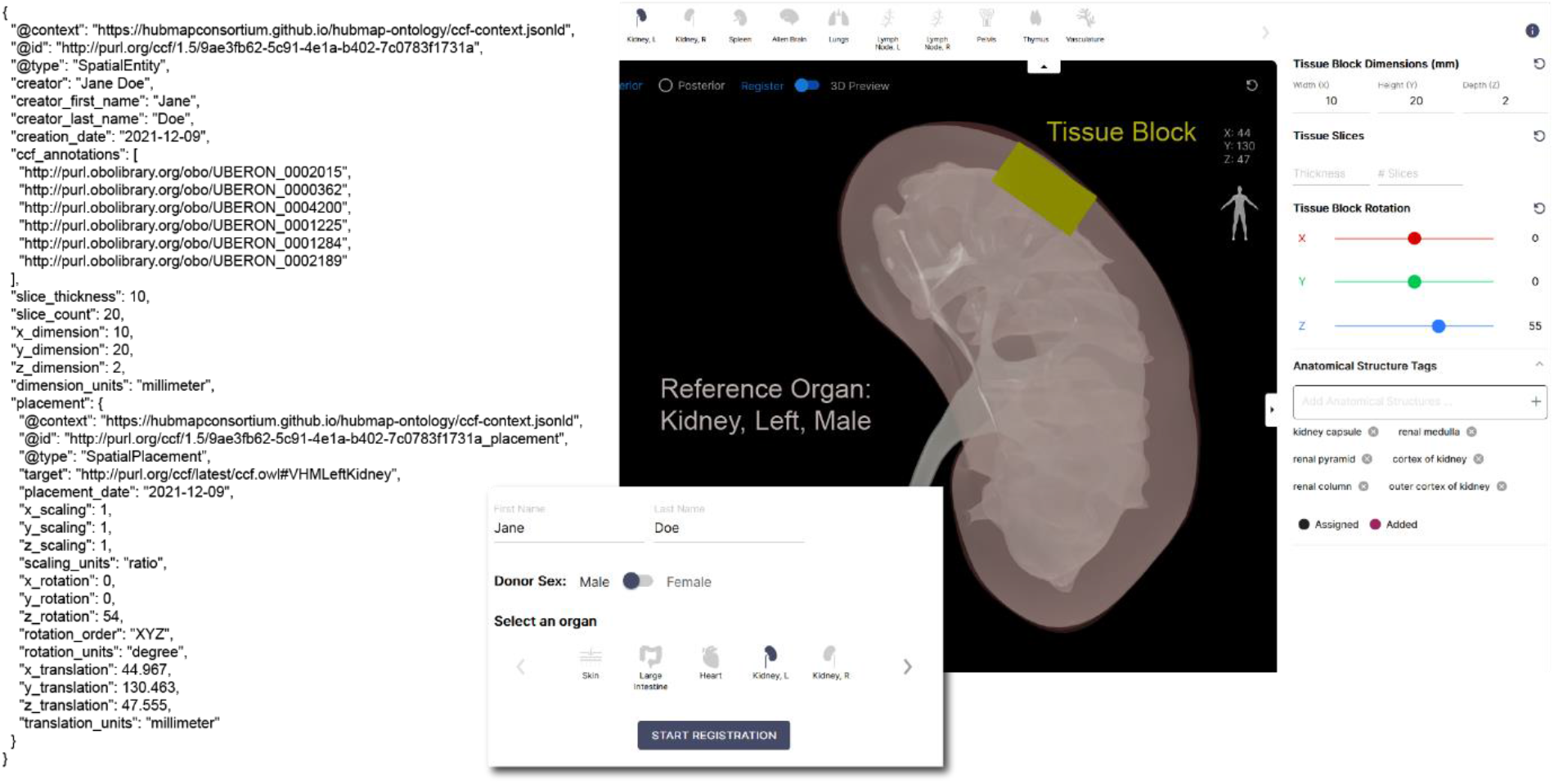
Registration User Interface Metadata. Exemplary JSON file with metadata for one tissue block registered in a kidney reference organ (the left male kidney) and RUI with user interface elements that capture all relevant information. The JSON file can be explored in the Supporting Information website at https://github.com/cns-iu/tissue-registration-and-exploration-UIs-in-support-of-hra/blob/main/rui_registration.json. All registered tissue blocks can be explored in the EUI.

**Supplementary Figure 4.**
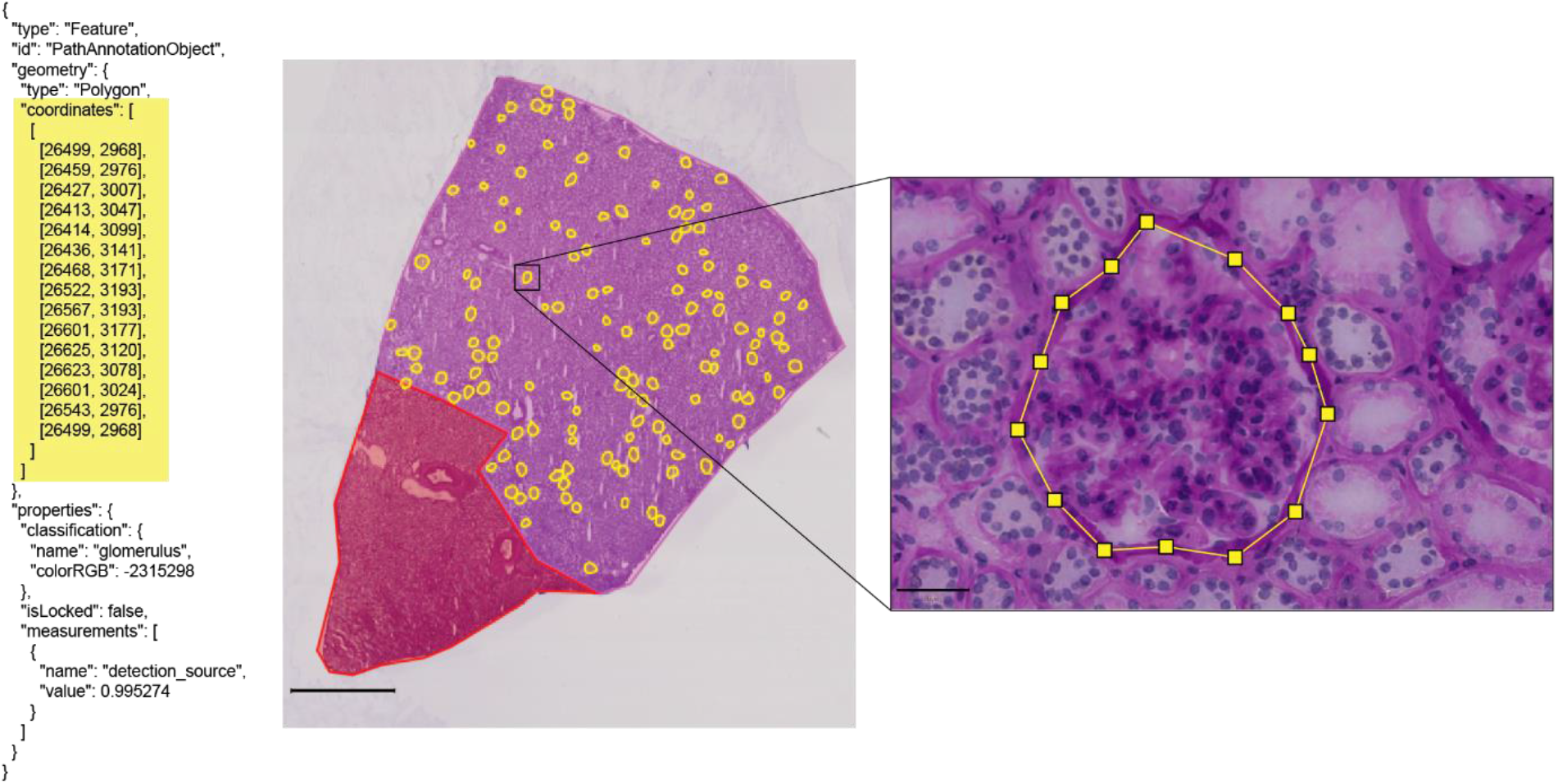
Kidney Tissue and Segmentation Data. Exemplary JSON file for a segmentation mask. Kidney whole slide image (scale bar: 2mm) with zoom into one glomerulus annotation consisting of 14-segment polyline, the associated 15 points are highlighted in yellow in JSON--note that the first set of coordinates is identical to the last closing the polyline (scale bar: 50µm).

## Notes

### Competing Interest Statement

The authors have declared no competing interest.

